# A motif preferred adenine base editor with minimal bystander and off-targets editing

**DOI:** 10.1101/2024.11.23.624961

**Authors:** Mengyu Shang, Yinuo Li, Qiuyu Cao, Jingxuan Ren, Yuqiang Zeng, Jinxin Wang, Xiyu Ge, Xiaohui Zhang

**Affiliations:** State Key Laboratory of Common Mechanisms Research for Major Diseases,Suzhou Institute of Systems Medicine, Chinese Academy of Medical Sciences & Peking Union Medical College, Suzhou 215123, China; Division of Endocrinology, Stanford University School of Medicine, Stanford, CA, USA

## Abstract

48% of hereditary disease are caused by single C-to-T base conversion, which makes efficient A-to-G base editing tools (ABEs) have great potential in the treatment of these diseases. However, the existing efficient ABE, while catalyzing A-to-G conversion, will bring more A and C bystander editing and off-target events, which poses safety concerns for their clinical application. Here, we developed ABE10 (ABE8e with TadA-8e A48E) for efficient and accurate editing of As in YA motifs with YAY>YAR (Y=T or C, R=A or G) hierarchy through structure-oriented rational design. Compared with ABE3.1, currently the only motif (YAC) preference ABE version, ABE10 exhibited A-to-G editing efficiency improvement with an average up to 3.1-fold in indicated YA motif while maintaining reduced bystander Cs editing and minimized DNA or RNA off-targets. Also, we showed ABE10 corrected pathogenic mutation with high efficiency and precision in human cells. Moreover, by ABE10, we efficiently and precisely generated hypocholes-terolemia and tail-loss mouse models mimicking human associated disease and mouse *PCSK9* base editing in vivo for hypercholesterolemia gene therapy, indicating their great potential in broad applications for and disease modeling and gene therapy.

## Introduction

About 60% of hereditary disease are caused by single base mutations, 48% of these are caused by single C-to-T base conversion^1^. The treatment of such hereditary diseases needs safe and effective genome editing tools. Traditional CRISPR/Cas9-mediated homologous recombination as tools to repair these pathogenic point mutations were very inefficient^1^, which leads to the new generation of gene editing tools-base editors, especially the A-to-G base editing tool (ABEs), providing an attractive platform for the treatment of single base mutation diseases^1, 2^.

Current popular base editors are cytidine base editors (CBEs) and adenine base editors (ABEs), which can convert C•G-to-T•A and A•T-to-G•C, respectively^2, 3^. They can be used to correct disease-causing point mutations that account for 48% and 13% of genetic diseases, respectively^1^. The application of gene editing in clinical treatment often requires accurate and efficient base editing tools. In recent years, scientists around the world have made significant progresses on the accuracy and efficiency of base editing. For CBEs, a series of accurate and effective CBEs have been developed, such as structure-oriented design of cytosine deaminase to construct motif-specific CBEs (T<u>C</u>, C<u>C</u> and WR<u>C</u>)^4–7^; Engineering cytosine base editors with only 1-2nt editing windows^8^; Re-evolving TadA-8e transformed ABEs into more accurate and less off-target CBEs^9–11^ etc. For ABEs, some progresses have also been made, such as rational design of accurate ABE9 for only editing A5 or A6 positions^13^. However, limitations of ABEs are still present for current ABEs, takingABE3.1 as an example, the only motif (YAC) preference ABE version, while has low efficiency and induced more bystander Cs editing within editing window and off-targets^2^. Therefore, ABEs that could efficiently catalyze A-to-G conversion in specific motifs without introducing more bystanders within editing window and off-targets still in urgent need.

Here, we obtained ABE10 with YA motif sequence preference through structure-oriented rational design of ABE8e mutant. Compared to ABE8e, ABE10 can precisely edit YA sequences with minimized bystanders. More importantly, compared with ABE3.1, the only motif (YAC) preference ABE version, ABE10 exhibited A-to-G editing efficiency improvement with average up to 3.1-fold in indicated motif while maintaining reduced bystander Cs editing and minimized DNA or RNA off-target. We also demonstrated that ABE10 can effectively correct disease-causing point mutations without introducing detrimental mutations in human cells. Moreover, by injecting ABE10 mRNA and sgRNA targeting splice donor of mouse *PCSK9* and *TBXT* into zygotes, the loss-of-function of PCSK9 mouse models for hypocholes-terolemia, and short-tailed mouse models for simulating mouse evolution were efficiently produced, respectively. Finally, we delivered lipid nanoparticle–packaged ABE10 mRNA and a sgRNA targeting the GT splice donor site of PCSK9 in mice, high base editing efficiency for PCSK9 and substantial reduction the expression of Pcsk9 and LDL-C in plasma were observed.

## Results

### Structure-based evolution of TadA-8e for YAC motif sequences preference

The earlier reported ABEs with YAC motif preference was ABE3.1^2^, but its A-to-G editing efficiency was low (1.2-31.1%) and induced more C bystander editing (Supplementary Fig. 1). Efficiency-wise, ABE8e is currently considered the more efficient version of ABE. However, ABE8e not only has higher editing activity and a wider editing window making it tend to induce more bystander edits and off-targets, but also lose its sequence preference^12^. To overcome this limitation, we designed a series of mutants of ABE8e at the binding and catalytic sites of TadA-8e as well as DNA substrates based on the crystal structure of ABE8e (Fig. 1a), and screened for ABEs to restore its YAC motif sequence preference while maintaining high editing activity. A human endogenous target site (HEK2) with A_3_C<u>A</u>_5_C<u>A</u>_7_A_8_ sequence was selected for testing. Among these mutations, ABEs that could edit A_5_ while had reducing or no editing efficiency of A_3_, A_7_ and A_8_ were selected as candidates (Fig. 1b). After co-transfected ABE8e mutants and HEK2 sgRNA for 72h in HEK293T cells, we collected genomic DNA of cells for amplicon high-throughput sequencing (HTS). HTS data showed that almost all ABE8e mutants can edit both A_5_ and A_7_, only ABE8e A48D exhibited reduced A_7_ base editing activity without sacrificing on-target A_5_ activity (Fig. 1c), indicating A48 maybe the key amino acid of TadA-8e that influences its motif sequence preference. In order to further identify the TadA-8e mutant that can minimize bystander editing, we saturated the mutant A48 amino acid and further tested it with the same target in HEK293T cells. HTS data showed that ABE8e A48E maintains comparable editing activity at A_5_ while minimizing bystander editing activity at A_7_ (a 3.0-fold decrease) compared to ABE8e A48D (Fig. 1d). We named ABE8e A48E as ABE10.

**Fig. 1.**
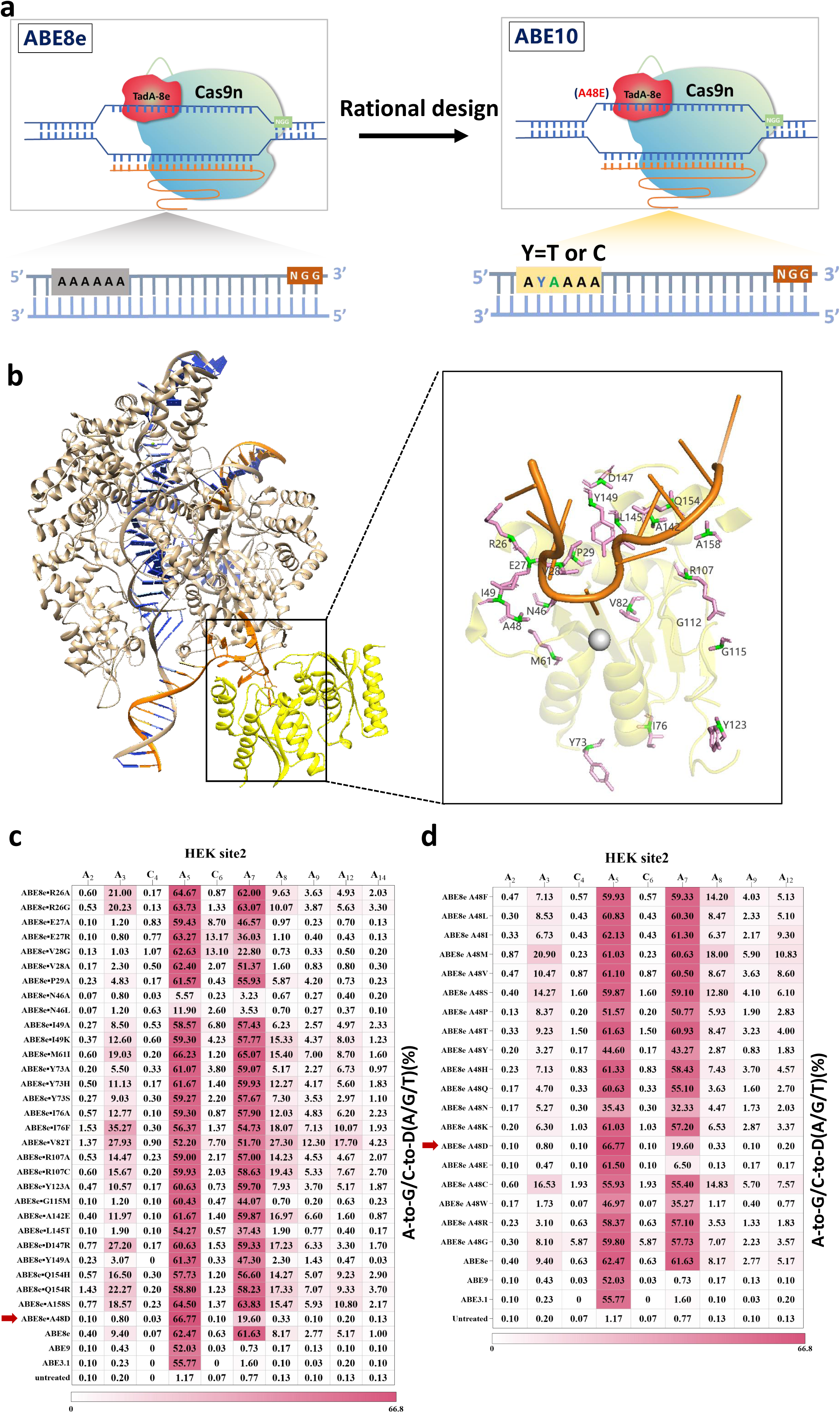
Rational design of motif preferred ABE. **a**, Chematic illustration of the conceptual design to design an efficient ABE into YA motif preferred ABE. **b**, Overview of the interaction of TadA-8e (yellow) with the single-stranded DNA substrate (orange) (Protein Data Bank (PDB): <u>6VPC</u>). Cas9n is in gray, sgRNA is in cyan, complementary strand DNA is in orange and noncomplementary strand DNA is in green. Amino acids spatially contacting or adjacent to the substrate DNA are labeled on the enlarged image. **c**, The A-to-G base editing efficiency of ABE8e or ABE8e variants at an endogenous genomic loci containing YAC motif and multiple As with non-YAC motif (HEK2) in HEK293T cells. Data are means (n = 3 independent experiments). **d**, The A-to-G base editing efficiency of ABE8e or ABE8e variants of A48 saturation mutagenesis at HEK2 in HEK293T cells. Data are means (n = 3 independent experiments).

### Characterization of ABE10

To unbiasedly profile the characteristics of ABE10, 23 endogenous targets were tested in HEK293T cells with ABE8e, ABE9 and ABE3.1 as control. Among these targets, they almost cover NAN (N=A, T, C and G) and some targets contained NCN (N=A, T, C and G) within its editing window. After analyzing the A-to-G editing efficiency of the most highly edited A in each target across all 23 target sites, we found that the efficiencies of ABE8e were 43.8-90.9%, which were significant higher than all other ABEs treated group (Fig. 2a,b). The efficiencies of ABE10 were 1.4-90%, whichwere significant higher than that of ABE9 (0.7-78.7%) and ABE3.1 (0-75.3%) (Fig. 2a,b). Through analyzing the A-to-G editing efficiency of each protospacer of all tested targets, we found ABE8e can efficiently edit almost all As without motif preference at major editing window (A_3_-A_8_) and even edit As outside of the major window (A_2_ or A_10_-A_13_) with considerable efficiency (Fig. 2a,c). ABE9, ABE3.1 and ABE10 have a similar editing window that was narrower than that of ABE8e, suggesting they are all accurate versions of ABE, suitable for precision gene therapy (Fig. 2a,c). We further analyzed the motif preference of these base editors and no motif preference for ABE8e and ABE9 was observed (Fig. 2d). Notably, ABE3.1 has YAY motif and ABE10 has YA motif preference with YAY>YAR hierarchy (Y=T or C, R=A or G) (Fig. 2a,d). Moreover, the average A-to-G efficiencies of ABE10 at A_4_, A_5_, A_6_ and A_7_ were 34.4%, 58.4%, 35.7% and 16.6%, respectively, far higher than the corresponding efficiencies of ABE3.1(Fig. 2a,c). Since ABE8e has been reported to have cytosine deaminase activity^14^, we further analyzed the C editing of ABE10 at 12 targets. The results showed that ABE10 induced lower or near background C-to-D editing activity compared to ABE8e. At *FANCF*-sg5, ABE10 decreased the editing efficiency of C induced by ABE8e by up to 10.4 times, which was comparable to ABE9 (Fig. 2e). ABE10 also induced no more indels than that of ABE8e and ABE3.1 (Fig. 2f). Together, ABE10 is highly efficient at YA motif preference with significantly reduced adenine and cytosine bystander editing effects.

**Fig. 2.**
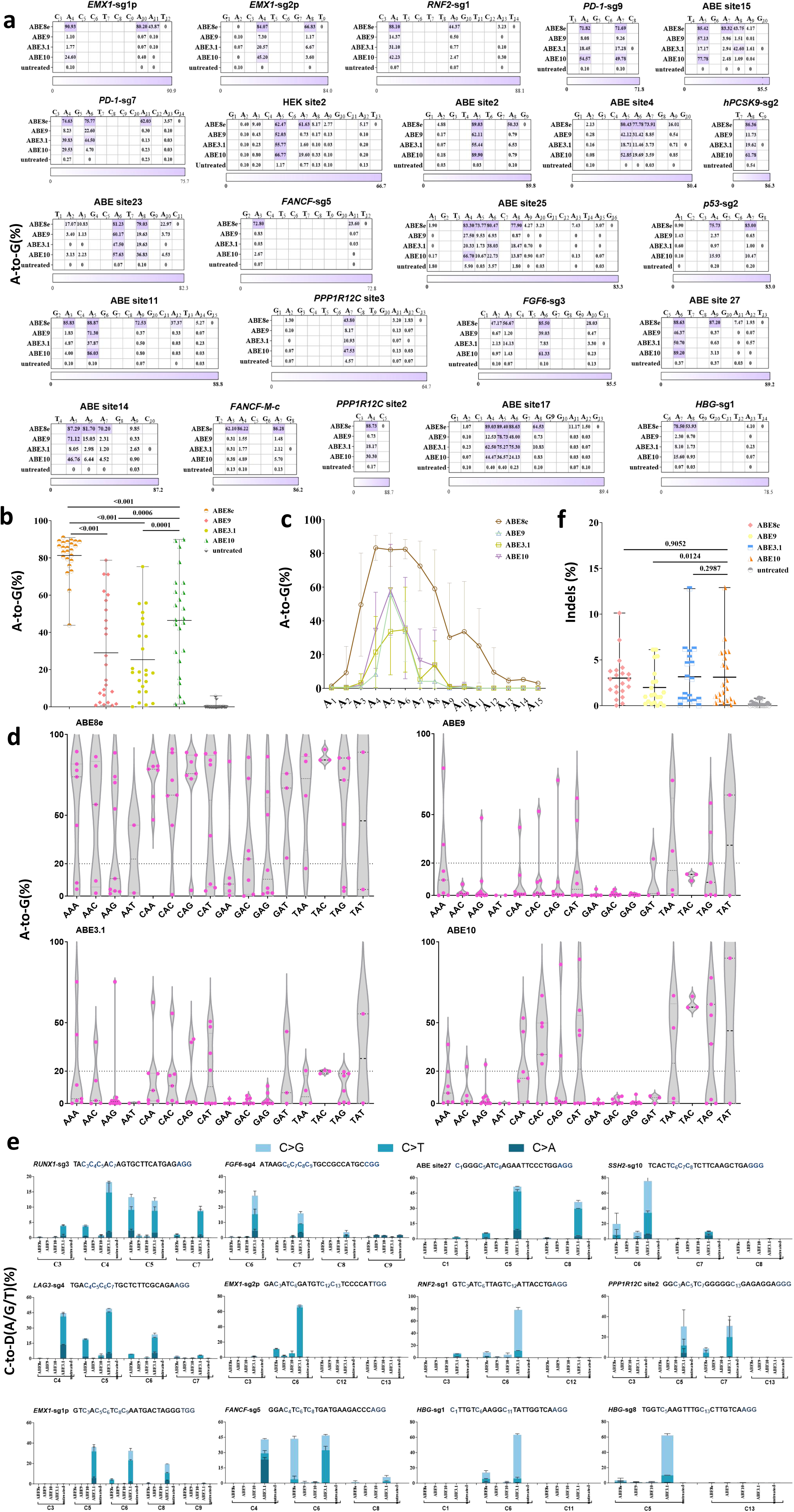
Characterization of ABE10. **a**, The A-to-G editing efficiency of ABE8e, ABE9, ABE3.1 and ABE10 were tested at 23 endogenous target sites in HEK293T cells. Data are means (n = 3 independent experiments). **b**, Summary of the A-to-G editing efficiencies for only the most highly edited adenine induced by ABE8e, ABE9, ABE3.1 and ABE10 were tested at 23 endogenous target sites in HEK293T cells. Each data point represents average A-to-G editing efficiency at all As within the activity window of each target site, calculated from 3 independent experiments. **c**, Average A-to-G editing efficiency of ABE8e, ABE9, ABE3.1 and ABE10 were tested at 23 endogenous target sites in HEK293T cells.Data are means ± SD (n = 3 independent experiments). **d**, The motif preferred analysis of ABE8e, ABE9, ABE3.1 and ABE10 at 23 endogenous target sites in HEK293T cells. Each data point represents average A-to-G editing efficiency at all motifs within the activity window of each target site, calculated from 3 independent experiments. **e**, Average C-to-D(A/C/G) editing efficiency of ABE8e, ABE9, ABE3.1 and ABE10 were tested at 23 endogenous target sites in HEK293T cells. Data are means ± SD (n = 3 independent experiments). **f**, The indels induced by ABE8e, ABE9, ABE3.1 and ABE10 at 23 endogenous target sites in HEK293T cells. Each data point represents average indels frequency at each target site calculated from 3 independent experiments. Significance was tested with two-sided paired Wilcoxon rank-sum test (**b**, **f**).

### Off-target evaluation of ABE10

Next, we performed off-target assessment of ABE10 in three ways: sgRNA-dependent DNA off-target, sgRNA-independent DNA off-target and whole-transcriptomic RNA off-target. 17 off-targets from in silicon prediction or Cas9 known off-targets were used to evaluate the sgRNA-dependent DNA off-target. The results showed that 4/17 off-target were observed for ABE8e and no sgRNA-dependent DNA off-target for both ABE10 and ABE9 (Fig. 3a). Modified R-loop assay was used to evaluate the sgRNA-independent DNA off-target^15^. HTS data showed that ABE10, similar to ABE9, induced no sgRNA-independent DNA off-target, which were far fewer than that ABE8e (Fig. 3b and Supplementary Fig. 3). RNA-Seq was used to evaluate whole-transcriptomic RNA off-target^16, 17^. The results showed that ABE10 induced 5 times lower the off-target rate than that of ABE8e, though slightly higher than that ABE9 (Fig. 3c). In summary, ABE10 is a highly efficient base editing tool with high specificity.

**Fig. 3.**
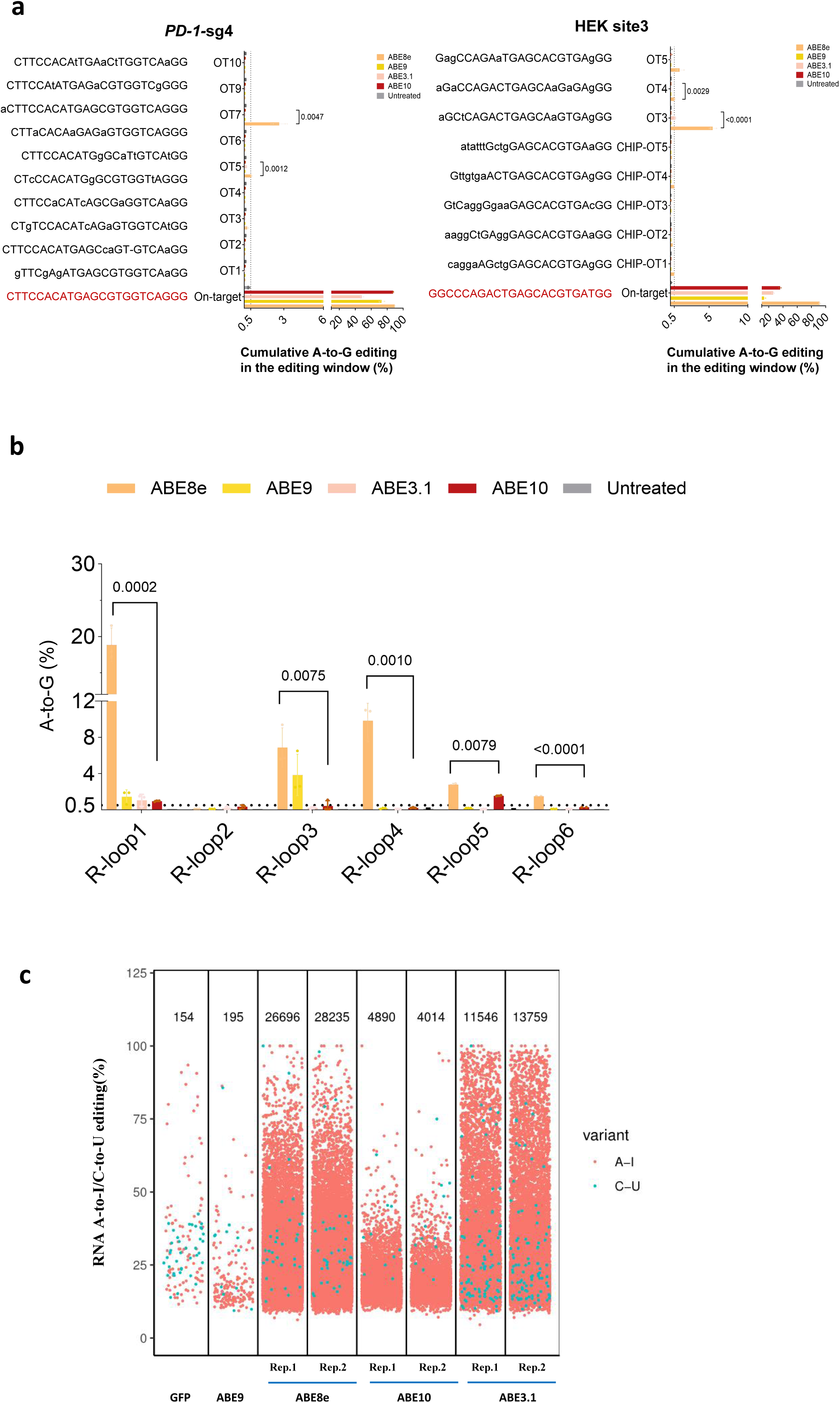
Off-target assessments of ABE10. **a**, Cas9-dependent DNA on and off-target analysis of the indicated targets (PD-1-sg4 and HEK3) by ABE8e, ABE9, ABE3.1 and ABE10 in HEK293T cells. Lowercase protospacer sequences represent mismatched bases compared to their corresponding on-target sequences. Data are means ± SD (n = 3 independent experiments). **b**, Cas9-independent DNA off-target analysis of ABE8e, ABE9, ABE3.1 and ABE10 using the modified orthogonal R-loop assay at each R-loop site with nSaCas9-sgRNA. Data are means ± SD (n = 3 independent experiments). **c**, RNA off-target editing activity by ABE8e, ABE9, ABE3.1 and ABE10 using RNA-seq. Each biological replicate is listed on the bottom. Significance was tested by two-tailed Student’s t test (**a**, **b**).

### Introduction of therapeutic mutation with high precision using ABE10 in human cells

To further validate ABE10’s potential for gene therapy with high precision and efficiency, we generated stable HEK293T cell lines with 2 pathogenic mutations (*GLAT* c.413C>T, Transferase Deficiency Galactosemia; *OTC* c.533C>T, Ornithine transcarbamylase deficiency) adjacent to bystander mutations that are deleterious in Clin Var database^18^. The results showed that ABE10 exhibited higher desired base editing efficiency than ABE9 at A_6_ in *GALT* (45.6%) and A_6_ in *OTC* (71.35%), though slightly lower than ABE8e at corresponding sites (Fig. 4a). However, ABE8e also induced deleterious no-desired base mutation with high efficiency, such as A_3_ in *GALT* and A_3_ in *OTC* which bring the possibility of additional disease. In contrast, ABE10 have far lower or no editing efficiency at these sites (Fig. 4a,b).

**Fig. 4.**
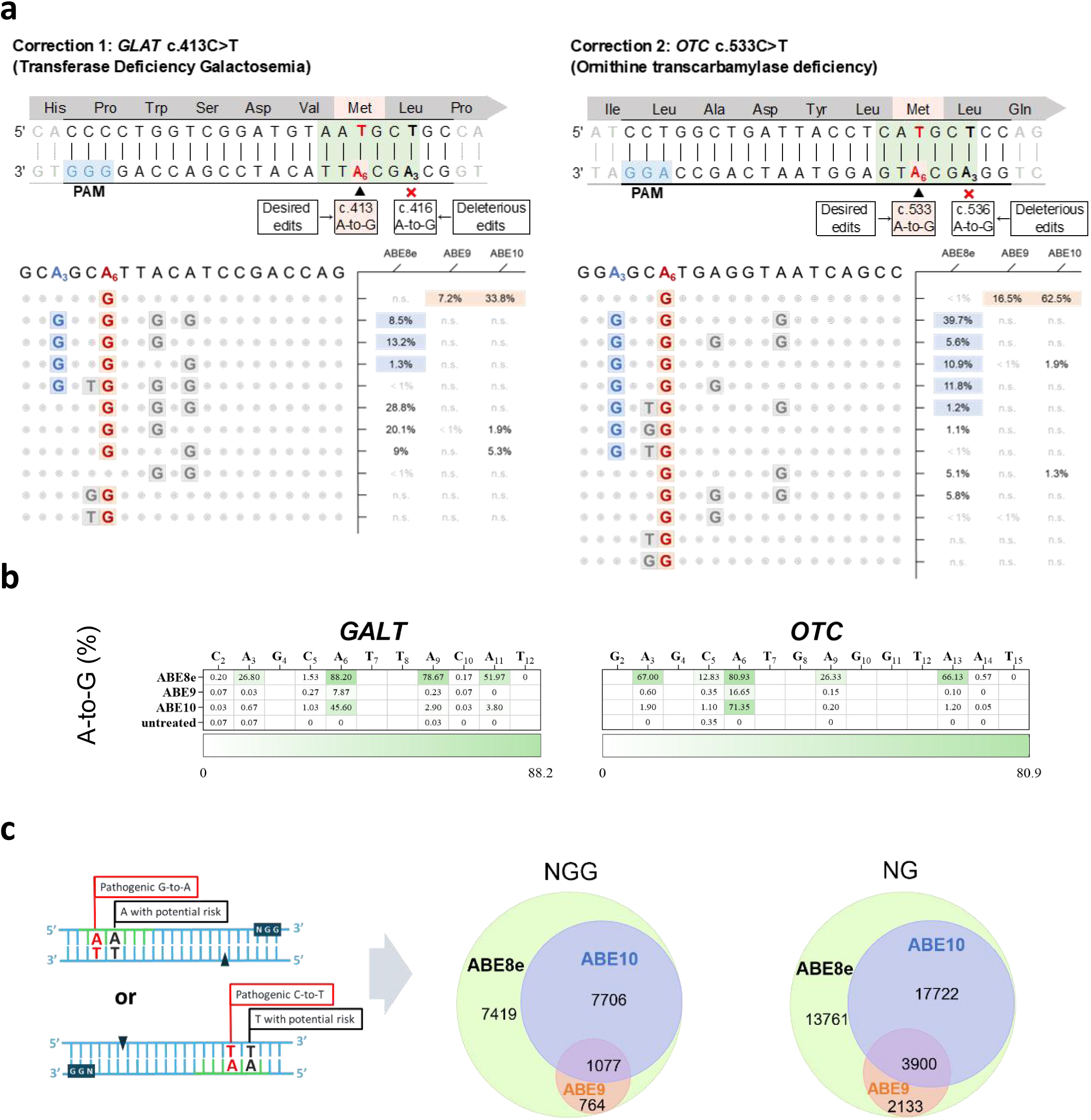
Pathogenic mutations corrected with high precision by ABE10. **a**, The allele base editing by ABE8e, ABE9 and ABE10 at HEK293T stable cells carrying 2 pathogenic mutations (*GLAT* c.413C>T, Transferase Deficiency Galactosemia; *OTC* c.533C>T, Ornithine transcarbamylase deficiency) adjacent to bystander mutations that are deleterious in Clin Var database. **b**, The A-to-G base editing of by ABE8e, ABE9 and ABE10 at the HEK293T stable cells. Data are means (n = 3 independent experiments). **c**, The statistics of pathogenic mutations corrected by ABE8e, ABE9 and ABE10 in Clin Var database (accessed July 16, 2024). The image on the left represents the schematic diagram of the precise correction of pathogenic mutations by ABEs, and the image on the right represents the total number of pathogenic mutations corrected by ABEs as PAM expands from NGG to NGN.

We further counted these pathogenic mutations with the presence of additional As or Ts (the disease-causing As or Ts were on the complementary strand of its DNA) that may present a risk of disease in the ClinVar database. The results showed that there were 12751 pathogenic mutations surrounded by As or Ts with potential security risk in ABE8ès editing window (Fig. 4c and Supplementary Table 4). Among those mutations, 8783 and 1841 pathogenic mutations could be precisely corrected by ABE10 and ABE9, respectively (Fig. 4c and Supplementary Table 4, 5). When PAM was further expanded from NGG to NG, the number of precise disease treatment events was further expanded to 21622 for ABE10 and 6033 for ABE9, respectively (Supplementary Table 4, 5). These data suggested that ABE10 have great potential in precisely targeting specific base for future clinical gene therapy.

### Generation of disease associated mouse models with high precision using ABE10

Base editors have been applied for making disease animal models that is essential for basic medical research. When the base editor is used to make the animal model of disease, it induces not only the target base conversion but also the bystander base editing which potentially perturb the analysis of the relationship between the mouse disease phenotype and base mutations^19^. Therefore, we used ABE10 to generate mouse model to minimize the occurrence of bystander events. We first targeted proprotein convertase subtilisin/kexin type 9 (PCSK9) to generate hypocholesterolemia mouse models by the zygotes microinjection of ABE10 mRNA and sgRNA targeting the splice site of PCSK9 (Fig. 5a and Supplementary Fig. 4)^20^. The results showed ABE 10 and ABE8e induced efficiently base editing in pups, with ABE 10(17/18) and ABE8e (9/11), respectively (Fig. 5b,c and Supplementary Fig. 4). However, only ABE10 induced precisely A-to-G conversion of the opposite DNA strand of GT splice site with minimized bystander editing in pups. On the contrary, not only the target base A_6_, but also bystander A_4_ was edited in ABE8e treated group (Fig. 5c and Supplementary Fig. 4). The phenotype of the pups carried precise target base editing by ABE10 were further confirmed by the detection of PCSK9 and LDL-C (Fig. 5d,e). These data suggest that ABE, especially the highly efficient version of ABE, enabled efficiently base editing in embryos to produce homozygous mouse with disease phenotype.

**Fig. 5.**
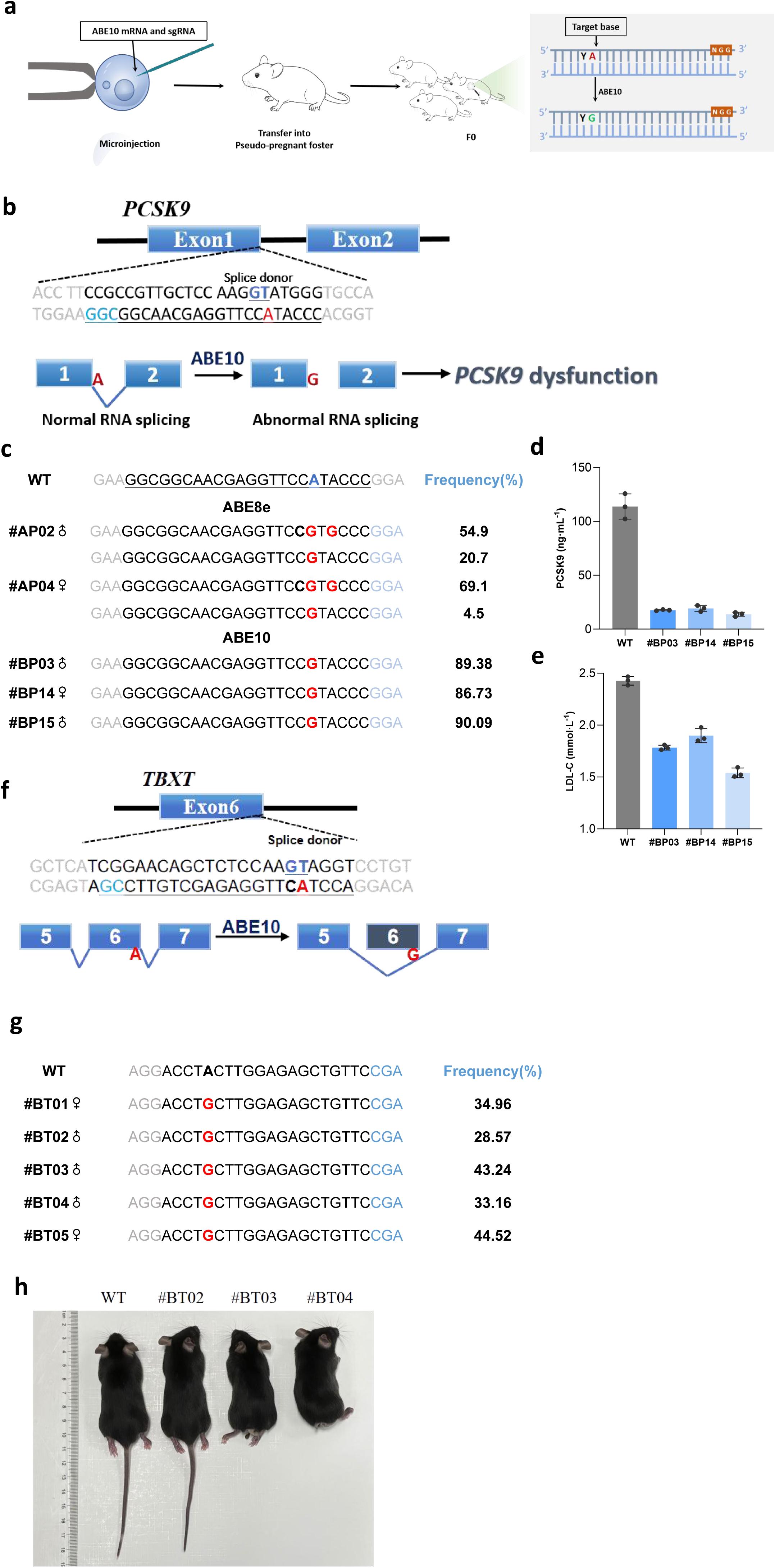
The generation of mouse models with high precision by ABE10. **a**, The workflow for generating mouse models by ABEs. **b**, The target sequence of ABE10 or ABE8e in the proprotein convertase subtilisin/kexin type 9 (*PCSK9*) exon 1 locus. The PAM sequence and sgRNA target sequence are shown in green and bold, respectively. The desired A-to-G conversion to disrupt the opposite DNA strand of the GT splice site is shown in red. **c**, Genotyping of representative F0 pups of *PCSK9* base editing by ABE8e and ABE10. The frequencies of wild-type (WT) and mutant alleles were determined by analyzing HTS using BE-Analyzer. The percentage values on the right represent the frequencies of the indicated mutant alleles. The frequency of the wild-type allele was omitted. **d**, **e**, The expression of PCSK9 and LDL-C in plasma of F0 pups from 8-week-old wild-type and founder (#BP03,#BP14 and #BP15) mice. Data are means ± SD (n = 3 independent experiments) **f**, The target sequence of ABE10 or ABE8e *TBXT* exon 6 locus. The PAM sequence and sgRNA target sequence are shown in green and bold, respectively. The desired A-to-G conversion to disrupt the opposite DNA strand of the GT splice site is shown in red. **g**, Genotyping of representative F0 pups of *TBXT* base editing by ABE10. The frequencies of wild-type (WT) and mutant alleles were determined by analyzing HTS using BE-Analyzer. The percentage values on the right represent the frequencies of the indicated mutant alleles. The frequency of the wild-type allele was omitted. **h**, The tail-loss phenotype of *TBXT* base editing F0 pups from 8-week-old wild-type and founder (#BT02,#BTO3 and #BT04) mice.

However, for some other disease-causing genes that are lethal with homozygous mutations, the highly efficient version of ABE is not suitable. For example, *TBXT*, a gene that influences the evolution of the mouse tail and single exon6-skipped transcript is sufficient to induce a tail-loss phenotype (Fig. 5f)^21^. We designed a sgRNA targeted the opposite of GT splice site of the *TBXT* that its disruption may possible cause exon6 skipping. By co-microinjection ABE8e or ABE10 with sgRNA into mouse zygotes, 8 (8/8) pups were edited in ABE10 treated group (Fig. 5g and Supplementary Fig. 5). Heterozygous mice (#BT03, #BT04) displayed almost no tail phenotype and mice with less editing efficiency (#BT02) showed short-tailed phenotype (Fig. 5h). However, we found no pups were born in ABE8e group (Data not shown) and the probable cause is that homozygous editing by ABE8e in mice leads to complete exon6-skipped and thus death in utero. These data suggest that ABE10 hold great potential in generation of disease model. More importantly, ABE10 has great advantages in animal models of diseases targeting homozygous lethal genes compared with other high efficiency version of ABEs.

### In vivo adenine base editing *PCSK9* in mouse using ABE10

To further validate ABE10’s potential for gene therapy in vivo, we targeted proprotein convertase subtilisin/kexin type 9 (*PCSK9*), a therapeutically relevant gene involved in cholesterol homeostasis^20^. We usedlipid nanoparticle–packaged delivery of mRNA encoding an ABE10 and a sgRNA targeting the GT splice donor site of *PCSK9* in mice (Fig. 6a). Three weeks after tail vein injection, higher base editing efficiency (60%) exclusively in the liver was observed (Fig. 6b), and no DNA off-target effects were detected by in silicon-predicated methods (Supplementary Fig.7a). During this time, we also monitored the expression levels of PCSK9 and LDL-C in plasma. The results also showed substantial reduction in the expression levels of Pcsk9 and LDL-C in ABE10 treated group (Fig. 6c,d). Meanwhile, LNPs-packaged ABE10 delivery did not induce liver injury by the detection of serum alanine aminotransferase (ALT) and aspartate transaminase (AST) (Supplementary Fig.7b,c). These data suggested that ABE10 has great potential in clinical gene therapy in vivo.

**Fig. 6.**
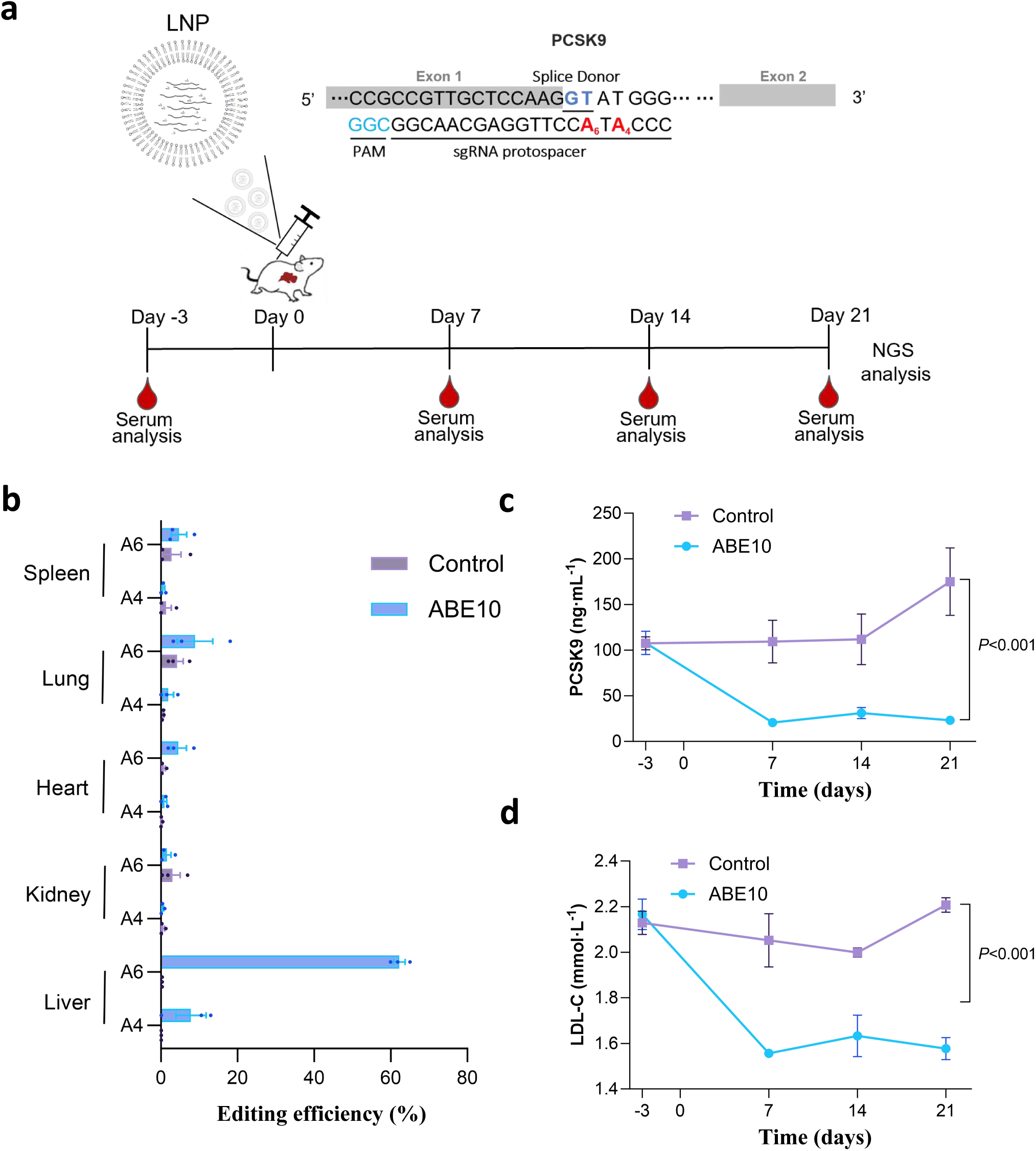
The mouse PCSK9 base editing in vivo using ABE10. **a**, The workflow for the mouse PCSK9 base editing in vivo by the delivery of LNPs packaged ABE10. The target sequence of ABE10 in the *PCSK9* exon 1 locus. The PAM sequence and sgRNA target sequence are shown in pink and blue, respectively. The desired A-to-G conversion to disrupt the opposite DNA strand of the GT splice site is shown in blod. **b**, The A-to-G editing efficiency of PCSK9 in adult mice 3-weeks after **t**he delivery of LNPs packaged ABE10 and sgRNA. Data are means ± SD (n = 3 different mice) **c**,**d**, The expression of PCSK9 and LDL-C in plasma of adult mice before and after **t**he delivery of LNPs packaged ABE10 and sgRNA. Data are means ± SD (n = 3 for the control group, n = 5 for the ABE10 group).

## Discussion

In this study, we have developed a YA motif preferred adenine base editor by structure-oriented rational design of TadA-8e and named it with ABE10. Compared to the efficient ABE version - ABE8e, ABE10 has YA motif preference and a narrower editing window, while reducing bystander A and C editing and DNA/RNA off-target. ABE3.1, currently the only sequence-specific ABE version, has unpredicatable C editing and RNA off-target. For ABE10, it not only effectively alleviates the limitations of ABE, but also has the advantages of high efficiency and small size. Compared with the current accurate ABE-ABE9, ABE10 not only has the same editing efficiency as ABE9 at A_5_ and A_6_, but also has higher editing efficiency at A_4_ and A_7_ sites, where ABE9 cannot edit. Therefore, ABE10 is an elegant adenine base editor with high efficiency, precision and small size. However, although ABE10 has high accuracy, it also sacrifices the editing efficiency of on-target. Therefore, it is necessary to further engineer deaminase in the future to improve its efficiency while maintaining its accuracy. For cytosine base editors, TC, CC, WRC motif preference cytosine base editors have been reported, while similar ABE tools are still lacking, and future further exploration is needed to enrich ABE’s toolbox.

Furthermore, we have also demonstrated that ABE10 can be effectively used to correct disease-causing point mutations in human cells, especially in editing windows where there is A mutation with safety risk. By further statistics in the ClinVar database, 7706 disease treatment events can be treated using ABE10 rather than other ABEs by precisely targeting YA motif. When PAM was further expanded from NGG to NG, the number of precise disease treatment events was further expanded to 17722, suggesting that ABE10 have great potential in precisely targeting specific base for future clinical gene therapy. Finally, ABE10 can be used to precisely target YA motif to generate animal models with disease phenotypes and to treat hypercholesterolemia without off-target events.

In summary, ABE10 is a highly efficient and specific base editing tool, which have great potential in animal model making and disease gene therapy.

## Methods

### Ethical statement

The research was conducted in accordance with all relevant ethical regulations. All procedures involving mice were meticulously designed to align with established guidelines and have received approval from the Institutional Animal Care and Use Committees (IACUCs) at the Suzhou Institute of Systems Medicine, Chinese Academy of Medical Sciences & Peking Union Medical College, Suzhou, China.

### Plasmid construction

The DNA sequences used in this research can be found in Supplementary Sequences. The ABE8e (#138489) and lentiCRISPR v2 (#52961) plasmids were obtained from Addgene. Polymerase chain reaction (PCR) was conducted using KOD-Plus-Neo DNA Polymerase (TOYOBO, Code: KOD-401). The plasmids generated in this article, including those based on ABE8e or lentiCRISPR v2 backbones, were constructed using the ClonExpress MultiS One Step Cloning Kit (Vazyme) (Supplementary Sequence). sgRNA expression plasmids were constructed as described previously^19^. Briefly, oligonucleotides listed in Supplementary Table 4 were denatured at 95°C for 5 min followed by slow cooling to room temperature. Annealed oligonucleotides were ligated into BbsI-linearized U6-sgRNA(sp)-EF1α-GFP for sgRNA expression (Thermo Fisher Scientific).

### Human Cell culture

HEK293T (ATCC CRL-3216) cell lines were cultivated in Dulbecco’s Modified Eagle’s medium (DMEM, Gibco) with 10% (vol/vol) fetal bovine serum (FBS, Gibco) and 1% Penicillin-Streptomycin (Gibco) antibiotic mix. HUDEP-2 cells were maintained and expanded in serum-free expansion medium (Stem Cell Technologies) supplemented with human Stem Cell Factor (SCF, 50 ng ml^−1^, PeproTech), erythropoietin (EPO, 3 IU ml^−1^, PeproTech), dexamethasone (DEX, 1 µM, Sigma), doxycycline (DOX, 1 µg ml^−1^, Takara Bio) and 2% penicillin–streptomycin (Gibco). All cell lines used were cultivated at 37 °C, 5% CO_2_ in the incubator.

### Stable cell line generation

The HEK293T stable cell line was constructed by cloning a 200-bp fragment containing disease-associated mutation flanked by ∼100 bp into lentiviral vector with puromycin-resistant gene expression maker. In brief, We used a combination of 12 μg of the specific lentiviral transfer plasmids (Lenti-GALT-sg1 and Lenti-OTC-sg1), along with 6 μg of pMD2.G and 9 μg of psPAX2, for co-transfection into HEK293T cells that were nearly 85% confluent in a 10-cm dish. The virus-containing supernatant was harvested at 72 hours post-transfection. The supernatant was subjected to centrifugation at 4,000 rpm for 10 minutes at 4 ℃, with the objective of precipitating cell debris. Following this, the supernatant was filtered through a 0.45 µm low protein binding membrane (Millipore). And then, it was serially diluted and added to a 24-well plate, each well containing 5 × 10^4^ HEK293T cells. After 24 hours, the lentivirus-transduced cells were spread into new plate wells supplemented with puromycin (2 µg ml^−1^). Following a week of puromycin selection, the pooled cells were spread into 96-well plate and single clone cells were harvested and expand for cell transfection.

### Cell transfection and fluorescence-activated cell sorting (FACS)

In DNA base editing experiments, HEK293T cells were plated in 24-well plates and transfected when they reached about 80% confluency. Next, a mixture of 3 μl polyethyleneimine (PEI, Polysciences), 1 μg plasmid DNA (750 ng ABEs or A&C-BEs expression plasmid and 250 ng sgRNA expression plasmid) and serum-free medium were applied to the cells. After three days, transfected cells were digested with 0.25% trypsin (Gibco) for fluorescence-activated cell sorting (FACS). Genomic DNA was isolated using the QuickExtract™ DNA Extraction Solution (QE09050, Epicentre) according to the manufacturer’s instructions. For RNA off-target analysis, HEK293T cells were seeded into 10-cm dishes and transfected with 30 μg of Cas9n-P2A-GFP, ABE8e-P2A-GFP, ABE9-P2A-GFP, ABE10-P2A-GFP and ABE3.1-P2A-GFP using PEI at approximately 80% confluency. After three days, transfected cells were washed with phosphate-buffered saline and digested with 0.25% trypsin (Gibco) for fluorescence-activated cell sorting (FACS). FACS was carried out on a FACSAriaIII (BD Biosciences) using FACSDiva version 8.0.2 (BD Biosciences). Cells were gated on their population via forward/sideward scatter after doublet exclusion (Supplementary Note). Then, cells (300,000 to 400,000 cells) with top 15% GFP signal were collected, and total mRNA was extracted using RNAiso Plus (Takara).

### Enhanced orthogonal R-loop assay

In this study, a modified orthogonal R-loop assay was employed for Cas9-independent DNA off-target analysis, which replaced dSaCas9-sgRNA plasmid with nSaCas9-sgRNA plasmid at each R-loop site. For the transfection process, a mixture of 3 μl polyethyleneimine (PEI, Polysciences), 850 ng plasmid DNA (250 ng SpCas9 sgRNA plasmid, 300 ng base editor plasmid (ABE8e, ABE9, ABE10, ABE3.1) and 300 ng nSaCas9 containing sgRNA plasmid were added to the cells. After three days, transfected cells were treated with 0.25% trypsin (Gibco) for sorting and then genomic DNA was extracted using the QuickExtract™ DNA Extraction Solution (QE09050, Epicentre), adhering to the manufacturer’s instructions.

### RNA sequencing (RNA-Seq) experiments

RNA sequencing was performed as previously described^19^. A total of 3 μg RNA per sample was used as input material for sample preparations. Sequencing libraries were prepared using a NEBNext Ultra RNA Library Prep Kit for Illumina (NEB) following the manufacturer’s recommendations. After assessing library quality with the Agilent Bioanalyzer 2100, index codes were added to assign sequences to individual samples. The index-coded samples were clustered on a cBot Cluster Generation System using the TruSeq PE Cluster Kit v3-cBot-HS (Illumina) according to the manufacturer ‘ s guidelines. Following cluster generation, the libraries were sequenced on an Illumina HiSeq platform, generating 125-bp or 150-bp paired-end reads.

### RNA sequence variant calling and quality control

RNA sequence variant calling and quality control were performed as previously described as follows^17^. Raw fastq format data were initially processed through in-house Perl Scripts. Adapter sequences were removed, and low-quality bases were trimmed to produce clean data using Trimmomatic. Concurrently, the Q20, Q30, and GC content of the clean data were calculated. All further analyses were conducted using this high-quality clean data. The reference genome index was built using HISAT2 version 2.0.5 and paired-end clean reads were aligned to the reference genome (Ensemble GRCh38) with the same HISAT2 version. Single-nucleotide polymorphism calling was carried out using GATK version 4.0. Variant loci in the base editor overexpression groups were filtered to eliminate sites without high-confidence reference genotype calls in the control group.

### High-throughput DNA sequencing and data analysis

On- and off-target genomic regions of interest were amplified via PCR using flanking high-throughput sequencing primer pairs listed in the Supplementary Table 2. PCR amplification was carried out with KOD-Plus-Neo DNA Polymerase according to the manufacturer’s instructions with 100-150 ng of genomic DNA as a template. For the preparation of high-throughput sequencing (HTS) libraries, site-specific primers containing an adaptor sequence (forward 5′-GGAGTGAGTACGGTGTGC-3′; backward 5′-GAGTTGGATGCTGGATGG-3′) at the 5′ end were employed in the PCR. Subsequently, the products were then subjected to a second-round PCR using primers with different barcode sequences. Then, PCR products with different tags were pooled together for deep sequencing using the Illumina HiSeq platform. For the batch analysis of the resulting FASTQ files, the reference sequences were set to full-length and analyzed as previously described^23.^ The base editing or indels efficiencies were quantified using using BE-Analyzer^23^ or CRISPResso^24^.

### Preparation of mRNAs and sgRNAs and Microinjection in mice

sgRNAs with the modification of phosphorothioation and methoxy group were synthesized by GenScript (Nanjing, China) (Supplementary Sequence). mRNA preparation was performed as previously described^19^. In brief, the T7 promoter was introduced into ABE8e or ABE10 template by PCR using the primers T7-ABEs-mRNA-F/R (Supplementary Table 2). The base editor mRNA were transcribed in vitro using mMESSAGE mMACHINE T7 Kit (Invitrogen) and purified using a MEGAclear Kit (Invitrogen)^20^.

C57BL/6J and ICR mice, housed in a specific pathogen-free environment with a 12-hour light/dark cycle and ad libitum diet, were used as embryo donors and foster mothers, respectively. A 2 nl mixture of ABE8e or ABE10 mRNA (200 ng μl^−1^) and sgRNA (100 ng μl^−1^) was co-injected into one-cell stage wild-type embryos. Injected zygotes were transferred into pseudopregnant female mice immediately using an Eppendorf TransferMan NK2 micromanipulator. About 20 days After embryo transplant, genomic DNA from born pups was isolated using the QuickExtract™ DNA Extraction Solution (QE09050, Epicenter) according to the manufacturer’s instructions.

### Animal Injection and Processing

All mice cohorts were maintained at specific pathogen-free (SPF) facilities in Suzhou Institute of Systems Medicine and approved by Institutional Animal Care and Use Committee. A total 200 μl mix of LNP packaged ABE10 mRNA and a sgRNA targeting the PCSK9 (40 μg: 20 μg) (Starna Therapeutics Co., Ltd., Suzhou) was delivered to 6-8 weeks old C57/BL6 mice intravenously via tail vein injection. A control group received an equivalent volume of normal saline. To track the serum levels of PCSK9 and Low-density lipoprotein cholesterol (LDL-C), mice were fasted overnight for 12 hours prior to blood collection via tail tip sampling. Blood samples were allowed to clot at room temperature, after which serum was separated by centrifugation and stored at -20°C for subsequent analysis. For terminal procedures, mice were euthanized using carbon dioxide inhalation. The median lobe of the liver was excised for DNA extraction to evaluate editing efficiency of on-target and off-target sites.

### Serum analysis

To minimize batch effects, serum samples from all time points were collected and analyzed concurrently. PCSK9 levels were measured using an ELISA kit (Proteintech, #KE10050), while LDL-C, ALT, and AST levels were assessed using assay kits from Solarbio (#BC5335, #BC1555, #BC1565, respectively). All procedures were conducted in strict adherence to the manufacturers’ instructions.

### Statistics and reproducibility

Three biologically independent replicates, conducted on different days, were used to calculate means and SD unless stated otherwise. All bar plots and figures were generated using Prism 9.3 (GraphPad). The statistical significance of differences between two groups was assessed using an unpaired two-tailed Student’ s t-test within Prism 9.3 (GraphPad). The specific P values can be found in the captions accompanying the figures. A P value of less than 0.05 was deemed to indicate significance. RNA-seq data was analyzed using Trim Galore (version 0.6.6), STAR (version 2.7.1a), SAMtools (version 1.14), Picard MarkDuplicates module (version 2.23.9) software. For the analysis of FACS data, FlowJo v.10 was employed.

## Data availability

The raw high-throughput sequencing data generated in this study have been submitted to the NCBI sequence Raed Archive database under SUB14831381, SUB14822433, SUB14822417. Additionally, the RNA-seq data are accessible in the NCBI sequence Read Archive database under accession code PRJNA818975. All plasmids described in this work will be deposited to Addgene. Source data are provided with this paper.

## Acknowledgments

We thank Professor Dali Li from East China Normal University for providing many base editors plasmids and some necessary experimental materials to accelerate the process of this project when I set up my research group. We thank Yuanqing He and Naiyun Ma from the Laboratory Animal Science Center of Suzhou Institute of Systems Medicine for their help in generating base-edited mice and the fund of NCTIB Cell and Gene Therapy R&D Platform. This work was partially supported by grants from National Key R&D Program of China (2022YFC3400203 to X.Z. and 2022YFA1103401 to X.Z.), the National Natural Science Foundation of China (No.32201223 to X.Z.), the Non-profit Central Research Institute Fund of Chinese Academy of Medical Sciences (2022-RC180-08 to X.Z.), the CAMS Innovation Fund for Medical Sciences (2022-I2M-1-024 to X.Z., and 2022-I2M-2-004 to Z.G.) and the Suzhou Municipal Key Laboratory (SZS2022005).

## Author contributions

M. S., Y.L., Q.C., J.R., J.W. and X.G. performed most of the experiments and analyzed the data. X.Z. designed and supervised the project. X.Z. wrote the manuscript with the input from all the authors.

## Competing interests

X.Z., H.L., J.R., Y.L. and Y.H. have submitted patent applications (application no. 202311786372.7, under review) based on the results reported in this study. This patent mainly relates to ABE10 in this paper. The remaining authors declare no competing interests.

## Notes

### Competing Interest Statement

The authors have declared no competing interest.

### Summary of Updates

Refine some of the details of Figure 1.

