## supplementary figures for "A motif preferred adenine base editor with minimal bystander and off-targets editing"

A-to-G/C-to-D(A/G/T)(%)

**_SSH2_-sg10**

| | $A_3$ | $C_4$ | $C_6$ | $C_7$ | $C_8$ |
|---|---|---|---|---|---|
| ABE8e | 56.93 | 0 | 49.63 | 0.93 | 0.43 |
| ABE3.1 | 1.17 | 3.93 | 87.60 | 9.23 | 0.97 |
| untreated | 0.03 | 0 | 0.07 | 0.03 | 0 |

Scale: 0 – 87.6

**_LAG3_-sg4**

| | $A_3$ | $C_4$ | $C_5$ | $C_6$ | $C_7$ |
|---|---|---|---|---|---|
| ABE8e | 79.33 | 0.43 | 19.57 | 4.87 | 2.40 |
| ABE3.1 | 13.57 | 44.67 | 48.97 | 23.77 | 3.40 |
| untreated | 0.10 | 0.17 | 0.10 | 0.10 | 0.40 |

Scale: 0 – 79.3

**HBG-sg1**

| | $C_6$ | $A_7$ | $A_8$ |
|---|---|---|---|
| ABE8e | 14.03 | 78.50 | 53.93 |
| ABE3.1 | 63.17 | 8.10 | 1.73 |
| untreated | 0 | 0.07 | 0.03 |

Scale: 0 – 78.5

**_HBG_-sg8**

| | $C_5$ | $A_6$ | $A_7$ |
|---|---|---|---|
| ABE8e | 3.70 | 41.60 | 37.37 |
| ABE3.1 | 62.30 | 10.00 | 6.93 |
| untreated | 0.03 | 0.07 | 0 |

Scale: 0 – 62.3

**RNF2-sg1**

| | $C_3$ | $A_4$ | $C_6$ | $A_9$ | $C_{12}$ |
|---|---|---|---|---|---|
| ABE8e | 0 | 88.10 | 8.70 | 44.37 | 0 |
| ABE3.1 | 7.50 | 31.10 | 78.07 | 0.77 | 0.50 |
| untreated | 0.10 | 0.07 | 0 | 0.07 | 0.10 |

Scale: 0 – 88.1

**EMX1-sg2p**

| | C3 | A4 | C6 | A8 |
|---|---|---|---|---|
| ABE8e | 0 | 84.07 | 11.03 | 66.83 |
| ABE3.1 | 1.90 | 20.57 | 68.33 | 6.67 |
| untreated | 0.10 | 0 | 0.17 | 0 |

Scale: 0 – 84.0

**Supplementary Fig.1 | High Cs editing induced by ABE3.1 in HEK293T cells.**
The A-to-G and C-to-D editing efficiency of ABE8e and ABE3.1 at 6 endogenous target sites in HEK293T cells. Data are means (n = 3 independent experiments).

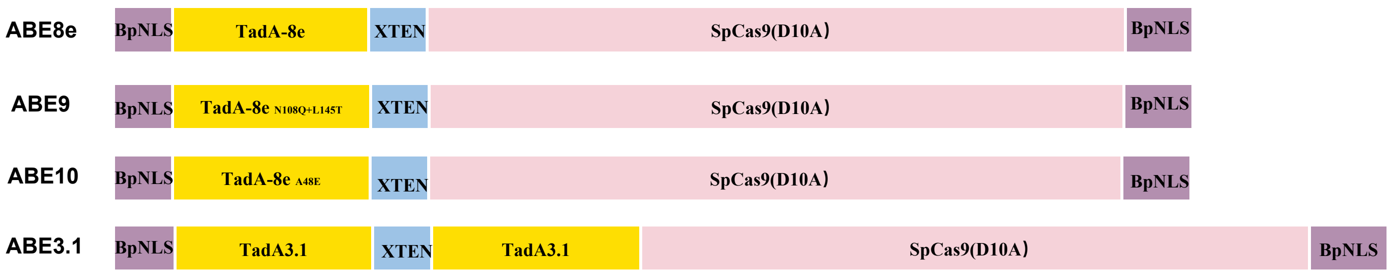


**Supplementary Fig.2 | Schematic representation of ABEs constructs.**
The bipartite nuclear localization signal (bpNLS) was in purple. The TadA and TadA variants was in yellow. The XTEN linker was in blue. The SpCas9 was in pink.

Supplementary Fig.3

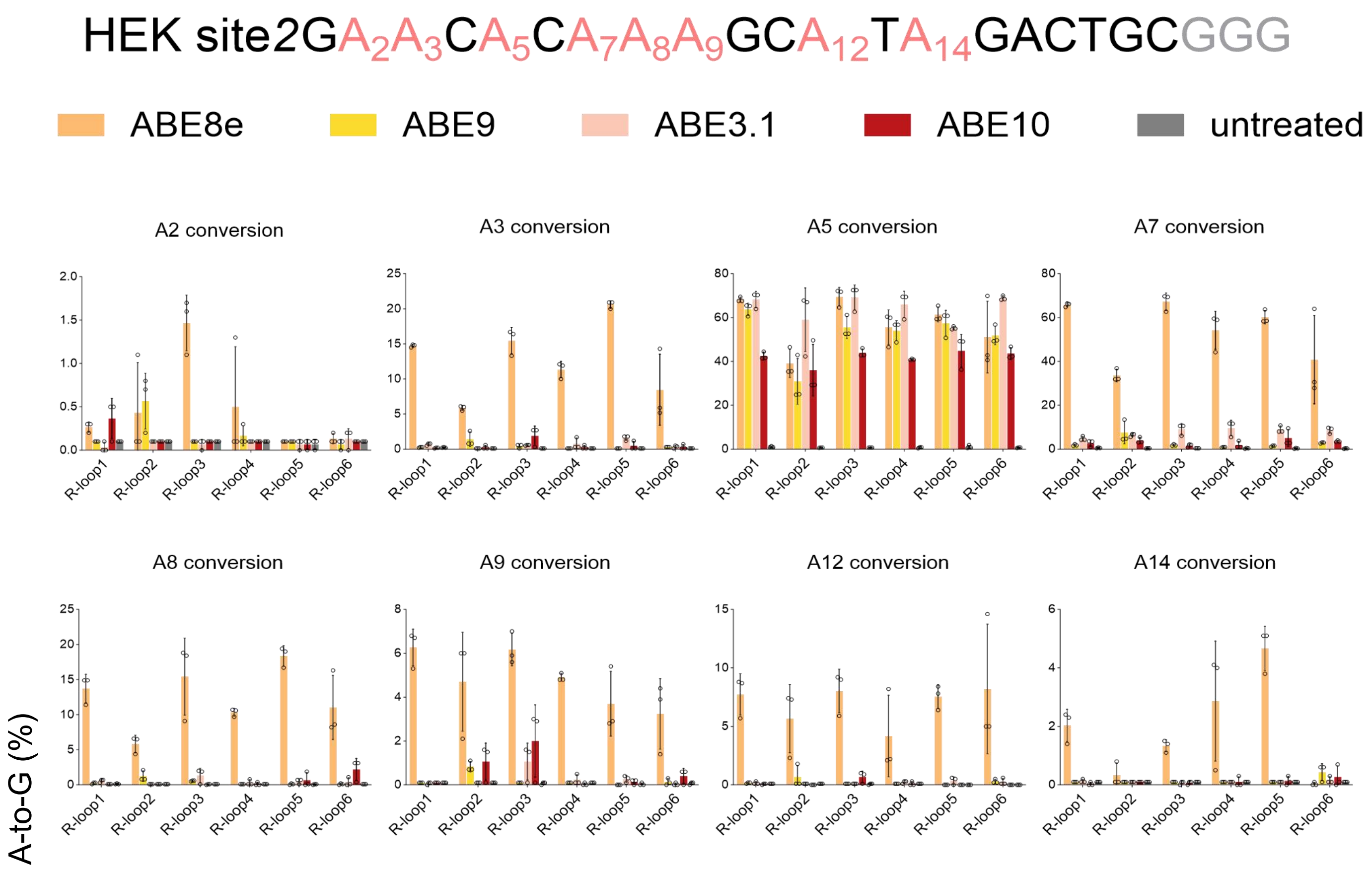


**Supplementary Fig.3 | The on-target A-to-G base editing efficiency by ABEs at HEK site2 in HEK293T cells.**
The on-target A-to-G base editing efficiency of ABE8e, ABE9, ABE3.1 and ABE10 at the endogenous genomic site (HEK site2) containing YA motif in Fig.3b. Data are means ± s.d. (n = 3 independent experiments).

**a**

| Gene | Base editor | No. of examined embryos | No. of transferred embryos (%) | No. of F0 pups (%) | F0 pups | |
|---|---|---|---|---|---|---|
| | | | | | No. of A-to-G mutants/total pups(%) | No. of Single A-to-G mutants/total pups(%) |
| PCSK9 | ABE8e | | | | | |
| PCSK9 | ABE10 | 193 | 81(42.0) | 18(22.2) | 17/18(94.4) | 16/18(88.9) |

**b**

| | Sequence | Frequency(%) |
|---|---|---|
| WT | GAAGGCGGCAACGAGGTTCCATACCCGGA | |
| | **ABE8e** | |
| #AP01 ♂ | GAAGGCGGCAACGAGGTTCCGTGCCCGGA | 待填 |
| | **ABE10** | |
| #BP01 ♀ | GAAGGCGGCAACGAGGTTCCGTACCCGGA | 89.02 |
| #BP02 ♂ | GAAGGCGGCAACGAGGTTCCGTACCCGGA | 88.85 |
| #BP04 ♂ | GAAGGCGGCAACGAGGTTCCGTACCCGGA | 88.47 |
| #BP05 ♂ | GAAGGCGGCAACGAGGTTCCGTACCCGGA | 89.78 |
| #BP06 ♂ | GAAGGCGGCAACGAGGTTCCGTACCCGGA | 24.85 |
| #BP07 ♂ | GAAGGCGGCAACGAGGTTCCGTACCCGGA | 83.38 |
| #BP08 ♀ | GAAGGCGGCAACGAGGTTCCGTACCCGGA | 89.52 |
| #BP09 ♂ | GAAGGCGGCAACGAGGTTCCGTACCCGGA | 89.85 |
| #BP10 ♂ | GAAGGCGGCAACGAGGTTCCGTACCCGGA | 89.77 |
| #BP11 ♂ | GAAGGCGGCAACGAGGTTCCGTACCCGGA | 46.62 |
| | GAAGGCGGCAACGAGGTTCCGTGCCCGGA | 43.78 |
| #BP13 ♀ | GAAGGCGGCAACGAGGTTCCGTACCCGGA | 89.36 |
| #BP16 ♂ | GAAGGCGGCAACGAGGTTCCGTACCCGGA | 90.26 |
| #BP17 ♂ | GAAGGCGGCAACGAGGTTCCGTACCCGGA | 90.50 |
| #BP18 ♂ | GAAGGCGGCAACGAGGTTCCGTACCCGGA | 90.40 |

**c**

Pcsk9 site

CCcTACCTTGGgGCAACaG
gaCATACCTTaaAGCAAaGG
aatgTACCTTGGAGCAACtG
ttCATACCTTGGAGCAAgGa
aCCATACCTTGGAaCAACca
CCCATACaTTGGgGCAtCGG
agCATACCaTGGAGCAACac
aCCATACCTaaGAGCAAact
agCATACCaTGGAGCAACac
CCCcTACCTTGGgGCAACaG
CCCATACCTTGGAGCAACGG

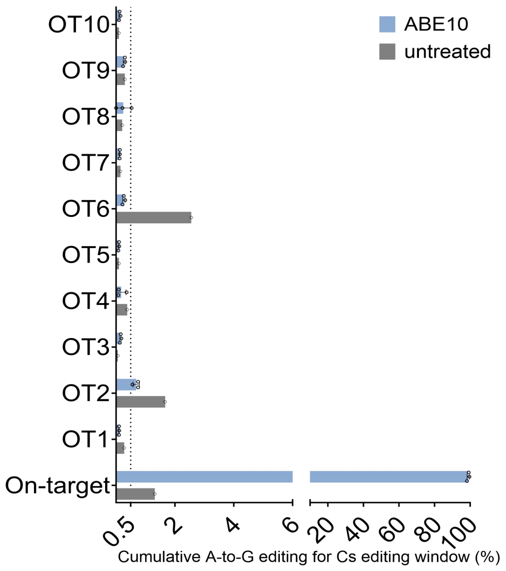


**Supplementary Fig.4 | High Precision base editing at the PCSK9 in Mouse Embryos by ABE10.**
a,The count of embryos utilized and the resulting pups following the microinjection of ABE8e and ABE10 mRNA/sgRNA.
b,Genotyping of representative F0 pups treated with ABE10 mRNA and sgRNA targeting the PCSK9 site. Wild-type pups (#BP12) were omitted. c, DNA on- and off-target analysis of the indicated targets PCSK9 site by ABE10 in F0 pups. Data are means ± s.d. (n= 3 independent experiments). Mismatched nucleotides in off-targeting sequences are indicated in lowercase.

**a**

| Gene | Base editor | No. of examined embryos | No. of transferred embryos (%) | No. of F0 pups (%) | F0 pups | |
|---|---|---|---|---|---|---|
| | | | | | No. of A-to-G mutants/total pups(%) | No. of Single A-to-G mutants/total pups(%) |
| *TBXT* | ABE8e | 139 | 90(64.7) | 0(0) | - | - |
| *TBXT* | ABE10 | 90 | 75/90(83.3) | 8(10.7) | 8/8(100) | 8/8(100) |

**b**

| | | Frequency(%) |
|---|---|---|
| WT | AGGACCT**A**CTTGGAGAGCTGTTCCGA | |
| #BT06 | AGGACCT**G**CTTGGAGAGCTGTTCCGA | 44.05 |
| #BT07 | AGGACCT**G**CTTGGAGAGCTGTTCCGA | 24.58 |
| #BT08 | AGGACCT**G**CTTGGAGAGCTGTTCCGA | 46.98 |

**c** ***TBXT* site**

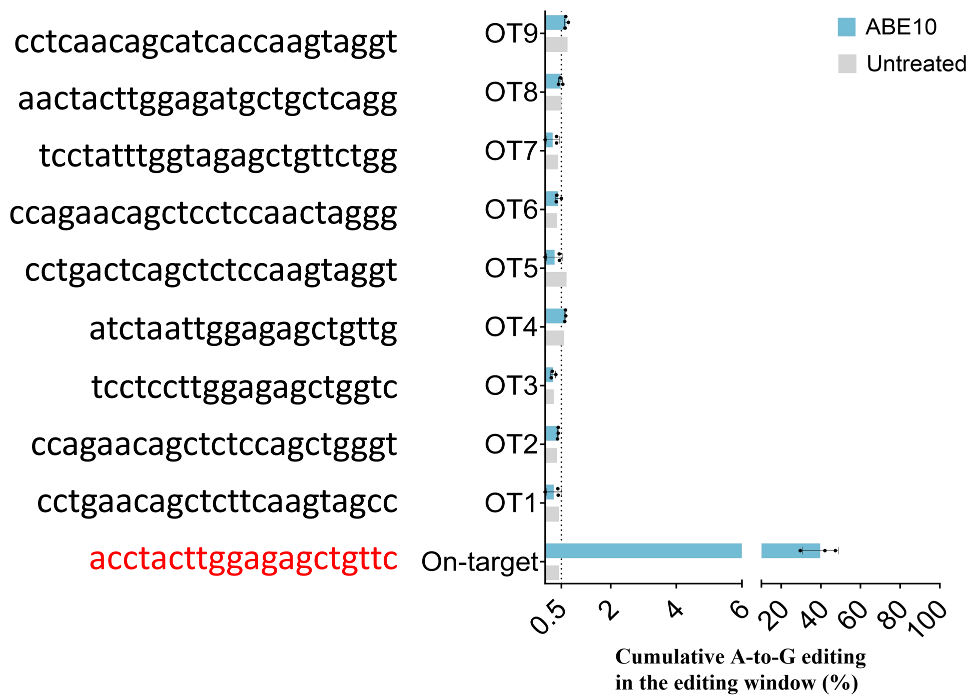


**Supplementary Fig.5 | High Precision base editing at the TBXT in mouse embryos by ABE10.**
a,The count of embryos utilized and the resulting pups following the microinjection of ABE8e and ABE10 mRNA/sgRNA. b, Genotyping of representative F0 pups treated with ABE10 mRNA and sgRNA targeting the TBXT site. c, DNA on- and off-target analysis of the indicated targets TBXT site by ABE10 in F0 pups. Data are means ± s.d. (n= 3 independent experiments). Mismatched nucleotides in off-targeting sequences are indicated in lowercase.

**Supplementary Fig.6**

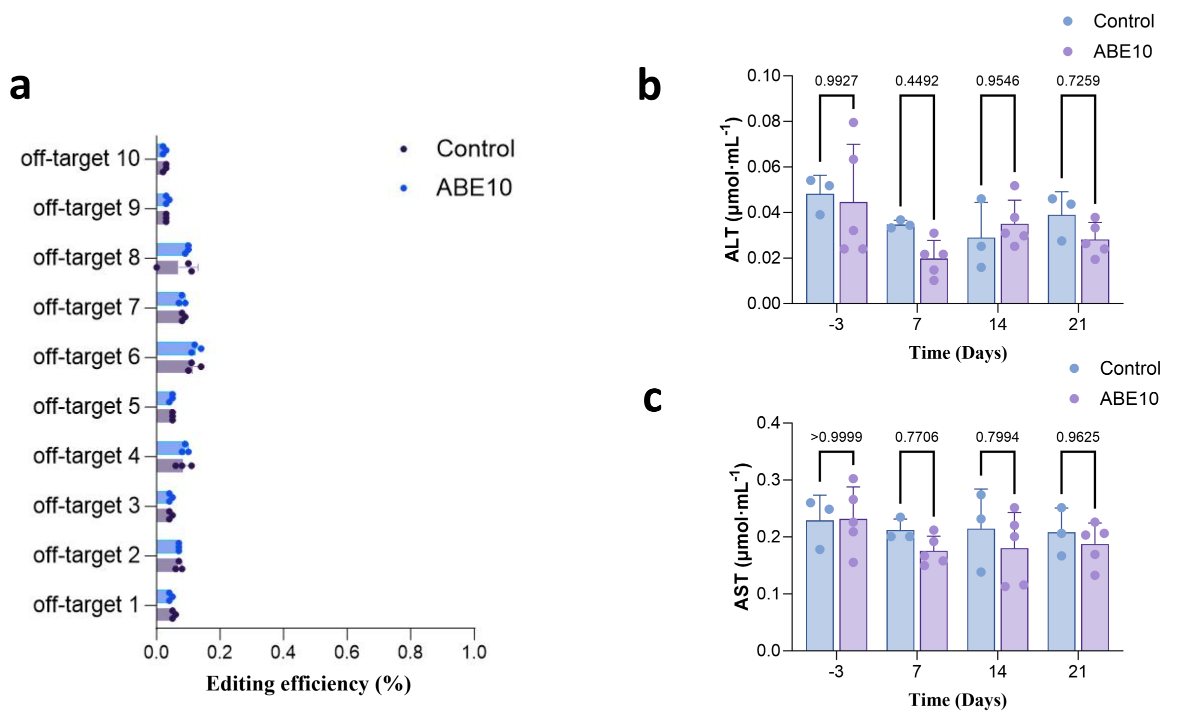


**Supplementary Fig.6 | Off-target editing efficiency and liver function following LNP Delivery of ABE10.**
a,Off-target editing efficiency measured across 10 potential off-target sites predicated by cas-offinder. The editing efficiency of ABE10 remains minimal and comparable to control. b, Alanine aminotransferase (ALT) and Aspartate aminotransferase (AST) levels in serum measured over time at days -3, 7, 14, and 21 following LNP delivery of ABE10 or control.

**HEK293T Negative Control（Untreated）**

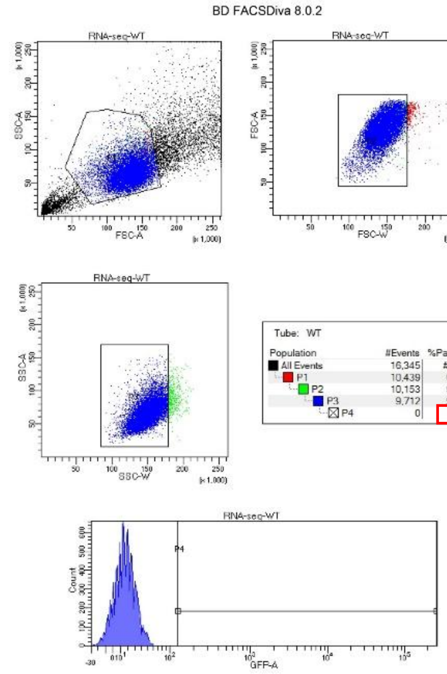


**HEK293T Positive Control（Cas9n-P2A-GFP）**

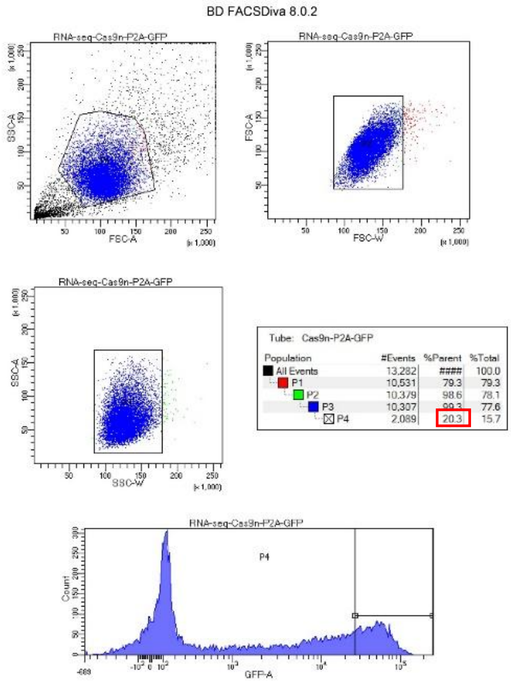


**ABE8e-P2A-GFP**

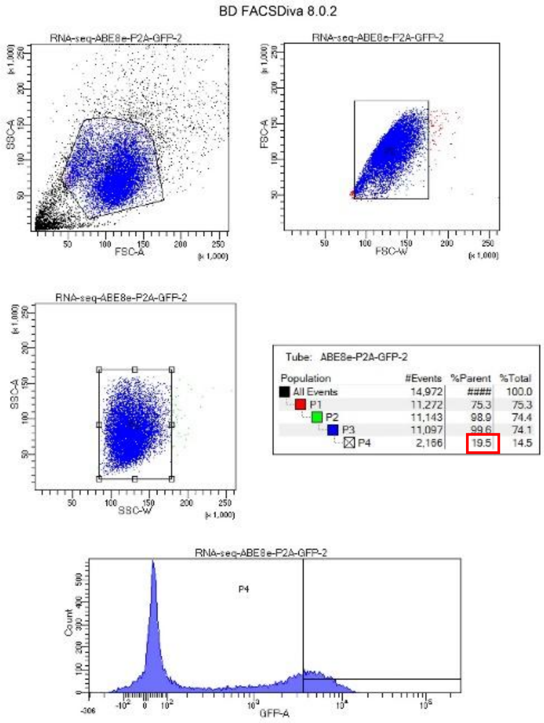


**ABE10-P2A-GFP**

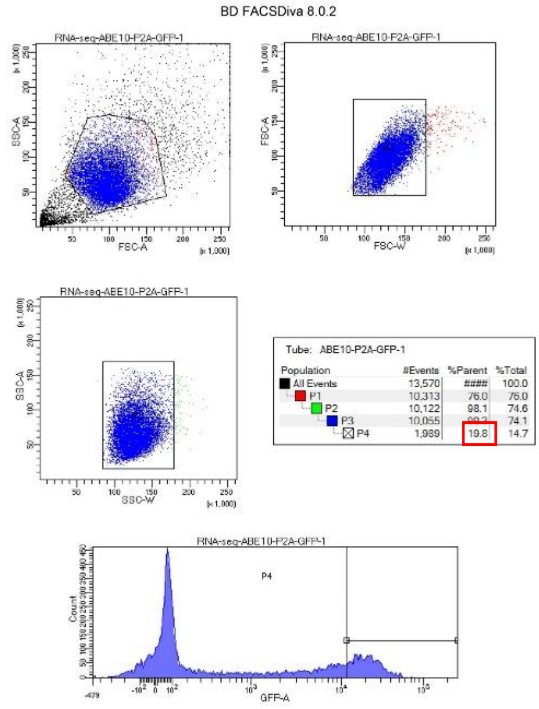


**ABE3.1-P2A-GFP**

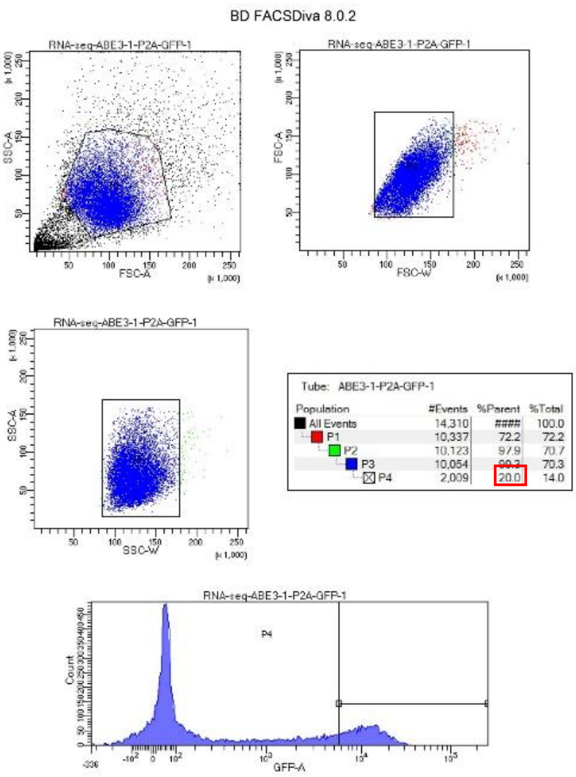


**ABE9-P2A-GFP**

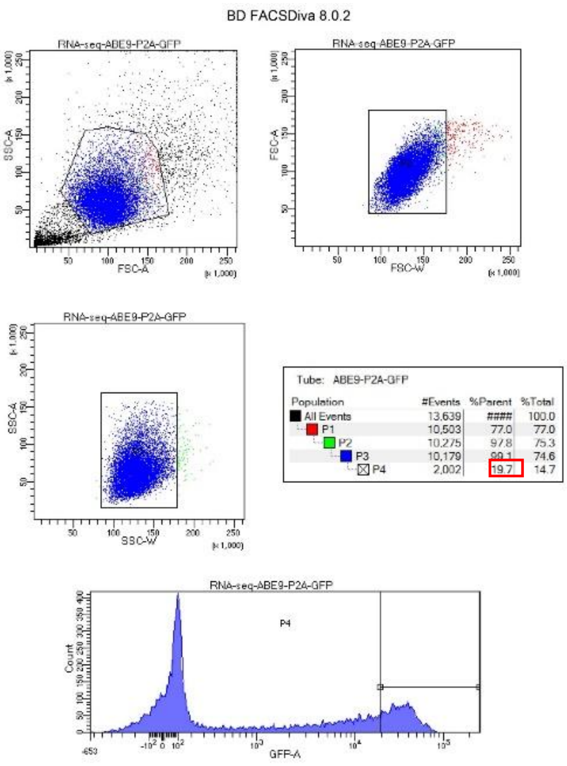


**Supplementary Note. FACS gating examples for GFP cell sorting conditions in transcriptome profiling.**
